# Disruption of ankyrin B and caveolin-1 interaction sites alters Na^+^,K^+^-ATPase lateral diffusion in HEK293 cell plasma membranes

**DOI:** 10.1101/141291

**Authors:** Cornelia Junghans, Vladana Vukojević, Neslihan N. Tavraz, Eugene G. Maksimov, Werner Zuschratter, Franz-Josef Schmitt, Thomas Friedrich

## Abstract

The Na^+^,K^+^-ATPase is a plasma membrane ion transporter of high physiological importance for ion homeostasis and cellular excitability in electrically active tissues. Mutations in the genes coding for Na^+^,K^+^-ATPase α-subunit isoforms lead to severe human pathologies including Familial Hemiplegic Migraine type 2 (FHM2), Alternating Hemiplegia of Childhood (AHC), Rapid Dystonia Parkinsonism (RDP) or epilepsy. Many of the reported mutations lead to change- or loss-of-function effects, whereas others do not alter the functional properties, but lead to e.g. reduced protein stability, reduced protein expression or defective plasma membrane targeting. Na^+^,K^+^-ATPase frequently assembles with other membrane transporters or cellular matrix proteins in specialized plasma membrane microdomains, but the effects of these interactions on targeting or protein mobility are elusive so far. Mutational disruption of established interaction motifs of the Na^+^,K^+^-ATPase with ankyrin B and caveolin-1 are expected to result in changes in plasma membrane targeting, changes of the localization pattern, and of the diffusion behavior of the enzyme. We studied the consequences of mutations in these binding sites by monitoring diffusion of eGFP-labeled Na^+^,K^+^-ATPase constructs in the plasma membrane of living HEK293T cells by fluorescence correlation spectroscopy (FCS) as well as fluorescence recovery after photobleaching (FRAP) or photoswitching (FRAS) and observed significant differences compared to the wild-type enzyme, with synergistic effects for combinations of interaction site mutations. These measurements expand the possibilities to study the consequences of Na^+^,K^+^-ATPase mutations and provide information about the interaction of Na^+^,K^+^-ATPase α_2_-isoform with cellular matrix proteins, the cytoskeleton or other membrane protein complexes.

## Introduction

The Na^+^,K^+^-ATPase belongs to the widely distributed class of P-type ATPases (Axelsen and Palmgren, 1998). It converts the energy from ATP hydrolysis to electrochemical gradients of Na^+^ and K^+^ across the cell membrane. The enzyme transports three Na^+^ ions out of and two K^+^ ions into the cell for each hydrolyzed ATP molecule. The resulting ion gradients are critical for the regulation of ion homeostasis, secondary active transport processes, for establishing the cellular resting potential, and for enabling action potentials during neuronal signaling or muscle contraction. The pivotal importance of Na^+^,K^+^-ATPase (or Na^+^ pump) is underlined by the fact that this enzyme can be the principal consumer of ATP in certain tissues, especially in the Central Nervous System (CNS), where the pump accounts for up to 70% of total ATP turnover.

The Na^+^,K^+^-ATPase is a hetero-oligomeric membrane protein, and its minimal functional unit comprises a large catalytic α-subunit with 10 transmembrane (TM) segments and a large cytoplasmic domain, and a smaller accessory β-subunit with one TM segment and a heavily glycosylated ectodomain (Figure 1). Depending on the tissue, another small single transmembrane-spanning γ-subunit is present (Figure 1), which belongs to the class of FXYD proteins (Sweadner and Rael, 2000). The α-subunit is responsible for the catalytic activity and ion transport of the enzyme and, as characteristic for P-type pumps, undergoes reversible phosphorylation coupled to distinct conformational transitions that coordinate ATP hydrolysis and cation transport. The β-subunit is required for the normal activity of the enzyme and for the structural and functional maturation of the α-subunit. Four α (α_1_, α_2_, α_3_, and α_4_) and three β-isoforms (β_1_ β_2_, and β_3_) have been identified in humans. Different isoforms combine to form Na^+^,K^+^-ATPase isozymes with distinct kinetic properties, which are regulated in a tissue- and developmental-specific manner. Within the same cell, more than one isoform can be expressed at the same time, and regulatory mechanisms have been developed by evolution to adjust the isoforms’ expression and activity to fulfill the particular physiological requirements (Friedrich et al., 2016).

**Figure 1:**
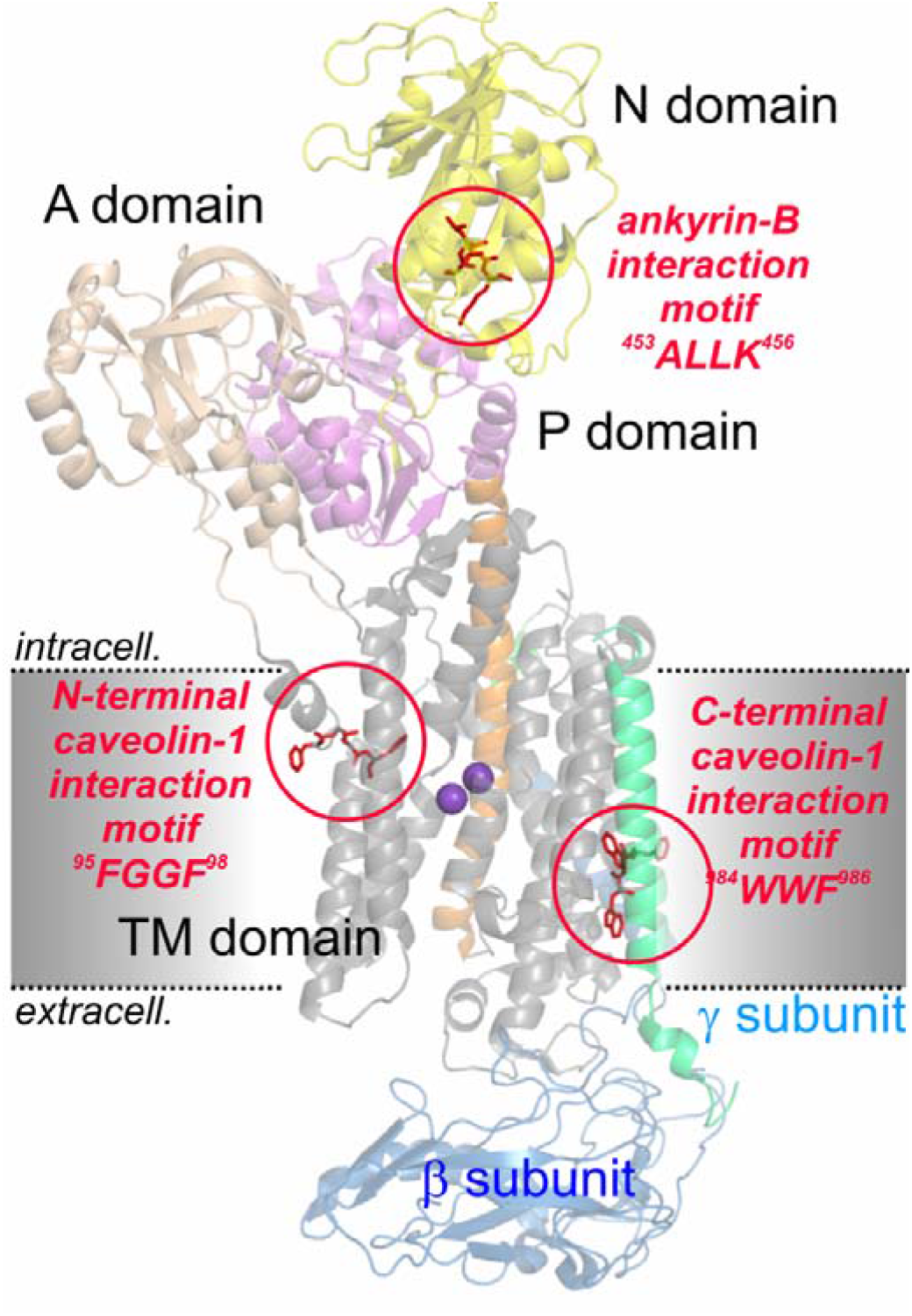
X-ray crystal structure of the Na^+^,K^+^-ATPase in Rb+-bound E_2_ conformation at 2.4 Å resolution (according to PDB entry 3A3Y) (Shinoda et al., 2009). Positions of the two interaction motifs with caveolin-1 (mutated in constructs NaK-ΔC, NaK-ΔN) and one for ankyrin B (mutated in construct NaK-K456E) are encircled. The functional domains of the Na^+^,K^+^-ATPase are indicated by different colors (N – nucleotide binding domain, yellow; P – phosphorylation domain, magenta; A – actuator domain, rose; transmembrane domain, grey, with M5 helix, orange). The β-subunit is shown in blue and the γ-subunit in cyan. Two violet spheres indicate bound Rb+ ions as congeners for K^+^.

Since 2003, genetic research has revealed that mutations in the human genes for α_2_ and α_3_ isoforms (*ATP1A2* and *ATP1A3*, respectively) cause severe neurological diseases including Familial Hemiplegic Migraine type 2 (FHM2) (De Fusco et al., 2003), Rapid Onset Dystonia Parkinsonism (RDP) (de Carvalho Aguiar et al., 2004), and Alternating Hemiplegia of Childhood (AHC) (Heinzen et al., 2012; Rosewich et al., 2012). The phenotypic spectrum of these distinct clinical entities overlaps significantly, and includes cognitive, movement and psychiatric disorders, migraines and epilepsies. This highlights the Na^+^,K^+^-ATPase as a paramount target for neurological disorders and has stimulated research on structure-function relationships, regulation and pharmacological interference of this important enzyme. A recent review about the involvement of the Na^+^,K^+^-ATPase α_2_-subunit in FHM2 and the effects of disease-related mutations can be found in (Friedrich et al., 2016).

Determining the pathophysiological mechanisms, by which the different functional outcomes of the mutations contribute to the pathogenesis of each of these diseases requires further structural and biochemical studies of mutant isoforms, as well as investigation of the Na^+^,K^+^-ATPase isoforms’ specific interactions with other cellular factors, and the development of appropriate disease model systems.

Whereas most mutations identified in FHM2 (and RDP or AHC as well) lead to distinct functional changes of the Na^+^,K^+^-ATPase that frequently support the notion of loss-of-function effects, several FHM2 mutations fail to exhibit functional alterations when assayed in a particular cell type (Tavraz et al., 2008). Thus, in order to completely address all possible reasons for pathophysiological consequences of Na^+^,K^+^-ATPase mutations in humans, functional studies must not only include electrophysiological and biochemical experiments in various relevant cell models, but should also assess effects regarding cellular quality control mechanisms, α/β subunit assembly, plasma membrane targeting, protein degradation, protein phosphorylation and - importantly - interactions with other proteins that might interfere with the aforementioned processes (Friedrich et al., 2016). Two major points on this list, which have not been scrutinized yet regarding their impact on pathogenesis, are the determinants of plasma membrane targeting and processes that are critical for the recruitment of Na^+^,K^+^-ATPase to distinct subdomains of the plasma membrane.

The impact of sub-cellular targeting of the Na^+^,K^+^-ATPase on its role in health and disease is still a matter of debate. It is known that Na^+^,K^+^-ATPase, in particular the α_2_-subunit found in glial cells (astrocytes) of the CNS or cardiomyocytes, forms complexes with the Na^+^, Ca^2+^ exchanger (NCX) and the inositol-1,4,5-triphosphate receptor (IP_3_R) in special Ca^2+^ signaling microdomains (Liu et al., 2008), or with NCX, the ryanodine receptor and the sarcoendoplasmic Ca^2+^-ATPase in so-called “junctional” endoplasmic reticulum (jER) microdomains, sometimes also dubbed PlasmERosomes (Blaustein et al., 1998; Juhaszova and Blaustein, 1997; Lencesova et al., 2004; Mohler et al., 2005; Stabach et al., 2008). Several reports indicated that the human Na^+^,K^+^-ATPase interacts with other cellular matrix proteins or constituents of the cytoskeleton to bring about sub-compartment localization. One such interaction partner for cellular targeting of the Na^+^,K^+^-ATPase is ankyrin B (Jordan et al., 1995; Liu et al., 2008; Zhang et al., 1998). Ankyrins are cytoskeletal proteins that interact via dynamic covalent bonds with integral membrane proteins and determine their distribution in the cell membrane. Three classes of ankyrins are distinguished, ankyrin R, ankyrin G and ankyrin B. The ankyrin B class is "broadly expressed" in most cell types and is encoded by the *ANK2* gene on human chromosome 4q25-27. All ankyrins are composed of four functional domains, an N-terminal membrane-binding domain consisting of 24 ankyrin repeats, a central domain that binds to the cytoskeletal protein spectrin, a death domain, as well as a C-terminal regulatory domain, which differs largely for the different ankyrin types. With its molecular weight of 440 kDa, ankyrin B (AnkB) is a particularly large representative and has an additional tail domain between the spectrin and the death domain consisting of 220 kDa of random coil sequence. Also, the membrane-binding domain differs from other proteins, which has specialized functions in unmyelinated axons and e.g. targeting of voltage-dependent channels. Ankyrin B and G are required for the polar distribution of many membrane proteins, including the Na^+^,K^+^-ATPase. The interaction with AnkB enables transport of the Na^+^,K^+^-ATPase from ER to Golgi and is associated with the stabilization of the enzyme in the plasma membrane. Importantly, mutations in the *ANK2* gene cause Long-QT syndrome type 4 (LQT4), a severe autosomal-dominant form of cardiac arrhythmia, by disrupting cellular organization of Na^+^,K^+^-ATPase (in particular of the α_2_-isoform that is located in transverse tubules), NCX, and IP_3_R, which are all AnkB binding proteins, in effect impairing the targeting of these proteins to transverse tubules as well as reducing the overall protein levels. One amino acid motif has been described in the literature (Jordan et al., 1995), by which the Na^+^,K^+^-ATPase α_2_-subunit interacts with AnkB. This Ala-Leu-Leu-Lys (ALLK) motif resides in the cytoplasmic nucleotide-binding (N) domain (Figure 1) comprising the positively charged lysine (Lys-456), which, upon mutation to a negatively charged glutamate, should drastically interfere with the AnkB interaction.

Another well-known Na^+^,K^+^-ATPase interaction partner is caveolin-1 (Cav1). Caveolins are structural elements forming the scaffold of caveolae membrane domains, building invaginations with a typical size of 50 to 100 nm in the plasma membrane e.g. of endothelial cells, which are visible in the electron microscope and even serve as protein markers for caveolae (Xie and Cai, 2003). So far, three different caveolin forms (caveolin-1, cavolin-2 and caveolin-3) with tissue-specific expression and distribution in mammalian cells are known that encode five different protein isoforms, from which Cav1 is the best characterized one. Cav1 has a molecular weight between 18 and 24 kDa and is found mostly in the plasma membrane (with two putative TM segments) and in Golgi-derived vesicles, with highest expression in fibroblasts, endothelial and smooth muscle cells. The protein forms high molecular weight complexes of about 13 to 16 monomers and is also interacting with caveolin-2, which is mostly co-expressed with cavolin-1, to form hetero-oligomers (Williams and Lisanti, 2004). Many cellular functions have been associated with caveolae and Cav1, such as membrane transport and endocytosis. They are also linked to the regulation of calcium and lipid metabolism, as well as to signal transduction during cell proliferation and programmed cell death. The Na^+^,K^+^-ATPase has been attributed to the regulation of caveolin trafficking and stabilization of Cav1 in the plasma membrane. Disruptions of caveolin targeting have been linked to several human pathologies such as dystrophies, disruption of cellular signaling cascades or endocytosis defects (Cai et al., 2008; Liu et al., 2002). Two Cav1 binding motifs on the Na^+^,K^+^-ATPase α-subunit have been described in the literature (Wang et al., 2004), which resemble the typical pattern φXXXXφXXφ or φXφXXXXφ (with φ representing an aromatic amino acid, X any amino acid). The N-terminal motif (**F**CRQL**F**GG**F**-98) resides at the intracellular interface of TM segment 2 and the C-terminal one (984-**W**W**F**CAFP**Y**) at the extracellular interface of TM segment 10 (see Figure 1). The consequences of mutational interference with these AnkB and Cav1 binding motifs are subject of this work, which aims to disclose whether interactions of the Na^+^,K^+^-ATPase α_2_-isoform with these proteins influence the cellular distribution pattern or the mobility of the enzyme in the plasma membrane.

Changes in the mobility of proteins in living cells can be probed by Fluorescence Correlation Spectroscopy (FCS), following the implementation of a specific fluorescence labeling scheme. FCS observes the fluorescence intensity fluctuations generated by molecules diffusing across a confocal volume of about 1 fL, and allows simultaneous observation of multiple diffusing molecular species without the need for their physical separation. To determine the diffusion behavior of different Na^+^,K^+^-ATPase constructs, N-terminal eGFP fusion proteins of human Na^+^,K^+^-ATPase α_2_-subunit were generated to enable the observation of Na^+^,K^+^-ATPase molecules for FCS studies. Different diffusion times and the diffusion coefficients calculated thereof provide information about the interaction of Na^+^,K^+^-ATPase α_2_-isoform with cellular matrix proteins, the cytoskeleton or other membrane protein complexes. The N-terminal Cav1 binding site was disrupted by mutations F95A/F98A (construct NaK-ΔN), the C-terminal Cav1 binding motif by mutations W984A/F986A (construct NaK-ΔC), and the AnkB binding motif by mutation K456E. Furthermore, two variants combining sets of these mutations (NaK-ΔNΔC and NaK-ΔNΔCK456E) were investigated to determine whether the effects of mutations are synergistic. As an alternative, we also employed fluorescence recovery after photobleaching (FRAP) and fluorescence recovery after photoswitching (FRAS) studies on some of these constructs and evaluated the use of the photoswitchable eGFP variant Dreiklang (DRK, (Brakemann et al., 2011)) for a comparison between FRAP or FRAS experimental schemes for discussion with results obtained by FCS.

## Materials and Methods

### cDNA constructs

The cDNA constructs of N-terminally eGFP-tagged (or Dreiklang-[DRK]-tagged) human Na^+^,K^+^-ATPase α2-subunit (harboring mutations Q116R and N127D in the ouabain binding region to confer reduced ouabain sensitivity, as reported in (Junghans et al., 2015) were generated within the pcDNA3.1X vector (Tavraz et al., 2008) by recombinant PCR. Three groups of amino acids within reported Cav1 or AnkB interaction motifs of the Na^+^,K^+^-ATPase α_2_-subunit were mutated using the QuickChange site directed mutagenesis kit (Stratagene), resulting in the following constructs: NaK-ΔN (disruption of N-terminal Cav1 binding site: mutations F95A/F98A), NaK-ΔC (disruption of C-terminal Cav1 binding motif: mutations W984A/F986A), NaK-K456E (disruption of AnkB binding motif: mutation K456E), as well as two constructs combining sets of these mutations: NaK-ΔNΔC and NaK-ΔNΔCK456E. All cDNA constructs were verified by sequencing (Eurofins MWG Operon, Ebersberg, Germany).

### Cell Culture and Transient Transfection

Human embryonic kidney (HEK293T) cells were cultivated at 37 °C in humidified 5 % CO_2_ atmosphere in DMEM cell culture medium with phenol red (Gibco), supplemented with 5 % fetal bovine serum and 1 % penicillin/streptomycin. For transient transfection, 300 μl of cells per well were seeded in eight-well Nunc™ Lab-Tek™ Chambered Coverglass with 1.0 borosilicate bottom (Thermo Scientific). 24 h after seeding, transient transfection was performed using Lipofectamine 2000 (Invitrogen) following the manufacturer’s protocol. 18 h to 24 h after transfection, the medium was replaced by DMEM Fluoro Brite (Gibco) with 10 % fetal bovine serum and 1 % penicillin/streptomycin. FCS measurements were performed 36 h to 48 h after transfection in a phenol red-free cell culture medium.

### Confocal Fluorescence Microscopy and FCS

Confocal imaging and FCS measurements were performed using a 510 META Laser Scanning Microscope with a ConforCor 3 system (Carl Zeiss, Jena, Germany). Excitation used the 488 nm line of an argon laser through a 40x water-immersion objective (C-Apochromat) with a numerical aperture (N.A.) of 1.2. The emitted light was collected through the objective and separated from the emission using a dichroic mirror (HFTKP 700/488). The resulting emission passed a 505 nm long pass filter before detection by an avalanche photodiode (APD). For imaging, the laser intensity was set to about 2 % of the 15 mW maximum laser intensity. For FCS measurements, the laser intensity was set to 0.5 % in order to avoid destruction of the sensitive biological material or photobleaching artifacts. FCS data were collected via a 70 μm pinhole in front of the APD detector. All experiments were carried out at 20°C.

Intracellular distribution of the fluorescence-labeled Na^+^,K^+^-ATPase constructs was visualized by confocal laser scanning microscopy. After cell imaging, a z-scan of fluorescence intensity was performed with 0.5 μm increments over a distance of 20 μm across the cell, and FCS experiments were carried out at the upper cell membrane identified by the respective peak in fluorescence intensity.

The autocorrelation curves (ACCs) were generated from 20 × 10 s-long data series. Only ACCs from cells exhibiting a stable count rate were taken for analysis. The measured autocorrelation curves G(τ) were normalized according to the following formula:

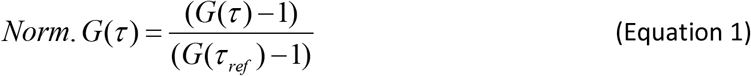

G(τ_Ref_) was calculated from the temporal average of the autocorrelation curve between *τ* = 5.2 · 10^−6^ s and 4.8 · 10^−5^ s. All normalized autocorrelation curves were plotted, and curves showing large deviations were discarded. Most of these deviating curves relate to bleaching processes or just showed no correlation.

Curve fitting used the integrated software of the Zeiss ConfoCor 3 system using the following model function for free two-dimensional diffusion with two components and a term for triplet transitions:

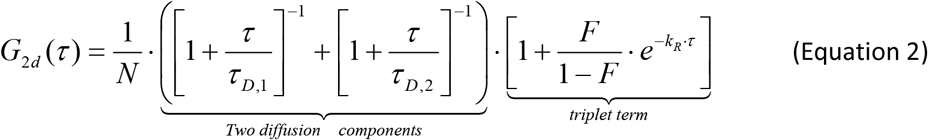

The beam waist r_0_ of the 488 nm laser line was calculated to be 0.17 μm based on reference measurements on rhodamine 6 G (R6G) with its known diffusion coefficient in aqueous solution (D = 2.8 · 10^−10^ m^2^·s^−1^ at 22 °C (Magde et al., 1974; Rigler et al., 1979)). The diffusion coefficients of the different eGFP-labeled Na^+^,K^+^-ATPase constructs were determined by using the calibrated beam waist *r*_0_ and the diffusion time τ_*D*_ according to the following equation:

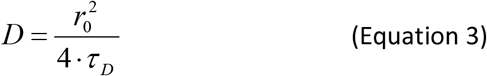

Two-sample Student’s t-test analysis was performed to determine whether the diffusion time components of the various Na^+^,K^+^-ATPase constructs were significantly different (P<0.05) from wildtype. Origin 9.1 was used for data analysis and presentation.

### Fluorescence Recovery after Photobleaching or Photoswitching

Confocal image series in fluorescence recovery after photobleaching (FRAP) experiments were acquired on a Leica TCS SP5 II microscope equipped with LAS AF acquisition software (Leica Microsystems; Wetzlar; Germany) using an 63x water-immersion objective with N.A. 1.4. For image series, a continuous-wave Argon laser was used and the laser intensity was set to 25 mW of the 30 mW maximum power in the focal plane. For cells expressing Dreiklang (DRK), the 514 nm Argon line was used for imaging (Intensity *I* = 8 % to 11 %) and bleaching (*I* = 100 %), and for eGFP-labeled samples, the 488 nm laser line was used (*I* = 6 % to 10 % for imaging, or 100 % for bleaching). As emission range for detection, 520 – 600 nm was used for DRK and 500 – 600 nm for eGFP. Fluorescence recovery after photoswitching (FRAS) measurements on DRK were performed using a 20× water-immersion objective with a N.A. of 1.0. In this case, a Coherent Cube 405 nm diode laser (Coherent, Santa Clara, USA) of the TCS SP5 II system (*I* = 100 %) was chosen to induce the OFF-switching process. All experiments were carried out at room temperature (22 °C).

FRAP and FRAS image series were recorded using the FRAP Wizard of the Leica TCS NT Software with an image size of 1024 × 128 pixels and 1400 Hz acquisition speed. First, 10 pre-bleach images were acquired (0.112 s time steps). Subsequently, bleaching (FRAP) or OFF-switching (FRAS) of a defined plasma membrane area was performed within a 5 ms bleach period at 100 % laser intensity followed by acquisition of 750 post-bleach images every 0.112 s (0.099 s for FRAS), followed by 100 images taken every 0.52 s using 2× line-averaging and bidirectional scanning.

FRAP/FRAS image sequences were analyzed using ImageJ software and the BioFormats plugin or the open source software easyFRAP (Rapsomaniki et al., 2012). Mean fluorescence intensities *I*(*t*) were calculated within a defined region of interest (ROI-1) representing the bleached membrane area. To compensate for bleaching effects over the scanning time, the averaged fluorescence intensity *I_ref_*(*t*) of a non-bleached reference membrane area (ROI-2) was determined. For background correction, the fluorescence intensity *I_bkg_*(*t*) outside the cell was determined (ROI-3). After background subtraction, a double normalization procedure was performed, which accounts for differences in starting intensities in ROI-1 and for the fluctuations of the total fluorescence signal during the experiment due to photobleaching or variations in the laser intensity:

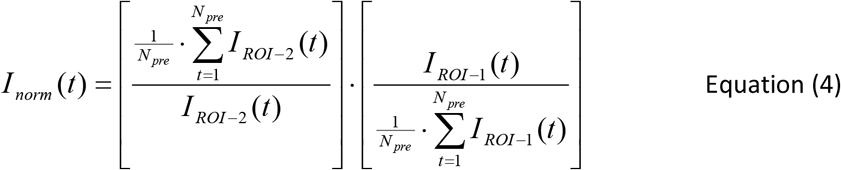

Here, the subscript “pre” symbolizes the pre-bleach image data and the time *t* refers to the number of the corresponding image. Subsequently, a full-scale normalization was performed according to:

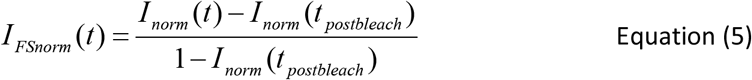

to additionally correct for differences in the bleaching depth.

Since the normalized intensity data resulting from this analysis were not well described with a single-exponential model function with one characteristic time, a bi-exponential function with two characteristic times was used for fitting using a non-linear least square minimization algorithm implemented in MatLAB:

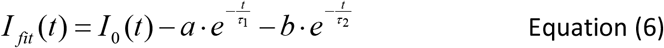

From these fit functions, the half-recovery time *t_1/2_*, defined as the time at which half-maximal recovery of the normalized fluorescence intensity from *I*_0_ to *I*_∞_ is being reached, was determined numerically for each individual cell. The half-recovery times (*t_1/2_*) were considered as diffusion times τ_*D*_ for the calculation of diffusion constants *D* according to the formula described in (Axelrod et al., 1976), which accounts for a small bleached spot in a two-dimensional area:

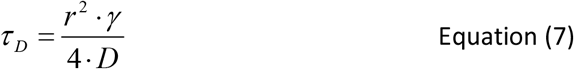

Here, *r* is the radius of the bleached spot, *γ* is introduced as a correction factor (set to 1 for full-scale normalization) for the amount of bleaching and τ_*D*_ is the diffusion time.

Since not all bleached molecules are mobile in the plasma membrane, fluorescence recovery (intensity at infinite time after bleaching, *I∞*) is usually incomplete, and the pre-bleach value of the fluorescence intensity *I_1_* will not be reached again. The difference of *I_1_ − I∞* is due to immobile molecules in ROI-1, which are not replaced by intact fluorescent molecules. Thus, the fraction

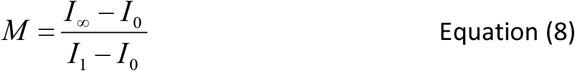

represents the mobile fraction of molecules in the observed area. Here, *I_∞_* indicates the normalized intensity in ROI-1 at infinite time of recovery, *I_1_* denotes the intensity before bleaching and *I_0_* just after bleaching. In case of a full scale normalization the formula reduces to *M = I_∞_*. More details about experimental methods and data analysis can be found in (Junghans, 2016; Junghans et al., 2015).

### Na^+^,K^+^-ATPase Expression in Xenopus oocytes

Ovary material was obtained by partial ovarectomy from anesthetized *Xenopus laevis* females, and individual cells were obtained by treatment with collagenase 1A (Sigma), as described (Tavraz et al., 2008). cRNA synthesis was carried out with the T7 mMessage mMachine kit (Ambion, Austin, TX). Each oocyte was injected with 25 ng of Na^+^,K^+^-ATPase α_2_-subunit and 2.5 ng of human Na^+^,K^+^-ATPase β_1_-subunit cRNAs and subsequently stored in ORI buffer (110 mM NaCl, 5 mM KCl, 1 mM MgCl_2_, 2 mM CaCl_2_, 5 mM HEPES, pH 7.4) containing 50 mg/L gentamycin at 18 °C for 3-4 days. Preceding the experiments, intracellular [Na^+^] was elevated by 45 min of incubation in Na^+^-loading solution (110 mM NaCl, 2.5 mM sodium citrate, 5 mM MOPS, 5mM Tris, pH 7.4) followed by an incubation of at least 30 min in Na buffer (100 mM NaCl, 1 mM CaCl_2_, 5 mM BaCl_2_, 5 mM NiCl_2_, and 2.5 mM MOPS, 2.5 mM Tris, pH 7.4).

### Rb^+^ Uptake Measurements

Uptake of Rb^+^ into oocytes was measured by atomic absorption spectrophotometry using an AAnalystTM 800 spectrophotometer (PerkinElmer Life Sciences, Rodgau, Germany) equipped with a Rubidium hollow cathode lamp (Photron, Melbourne, Australia) and a transversely heated graphite furnace. After Na^+^ loading, the oocytes were incubated for 3 min at 21–22 °C in Na buffer supplemented with 1mM RbCl and 10 μM ouabain to inhibit the endogenous Na^+^,K^+^-ATPase of oocytes. After three washes in Rb+-free Na buffer and one in Millipore water, individual cells were homogenized in 1 ml Millipore water. From these homogenates, 20 μl samples were injected into the graphite furnace for analysis using the autosampler of the AAS instrument. More details can be found in (Dürr et al., 2013; Friedrich, 2015).

## Results

To clarify whether the mobility of the Na^+^,K^+^-ATPase in the plasma membrane of cells is dependent on known interactions with cellular matrix proteins such as AnkB or Cav1, mutant Na^+^,K^+^-ATPase constructs carrying an N-terminal eGFP label were generated and the diffusion behavior in the plasma membrane of HEK293T cells was analyzed by FCS.

### FCS experiments

Figure 2 shows confocal scanning fluorescence microscopy (CSFM) images of cells expressing the various Na^+^,K^+^-ATPase constructs. Compared to the cytoplasmic distribution pattern of eGFP fluorescence, all Na^+^,K^+^-ATPase constructs exhibited profound plasma membrane expression, as expected. Whereas the wild-type construct shows the most prominent fluorescence in the plasma membrane, in particular the fluorescence signals for NaK-K456E, the double mutant NaK-ΔCΔN and the triple mutant NaK-ΔCΔNK456E exhibited a more diffuse pattern, with less fluorescence signal from the plasma membrane compared to intracellular fluorescence staining indicating that transport to the plasma membrane is already affected by these mutations.

**Figure 2:**
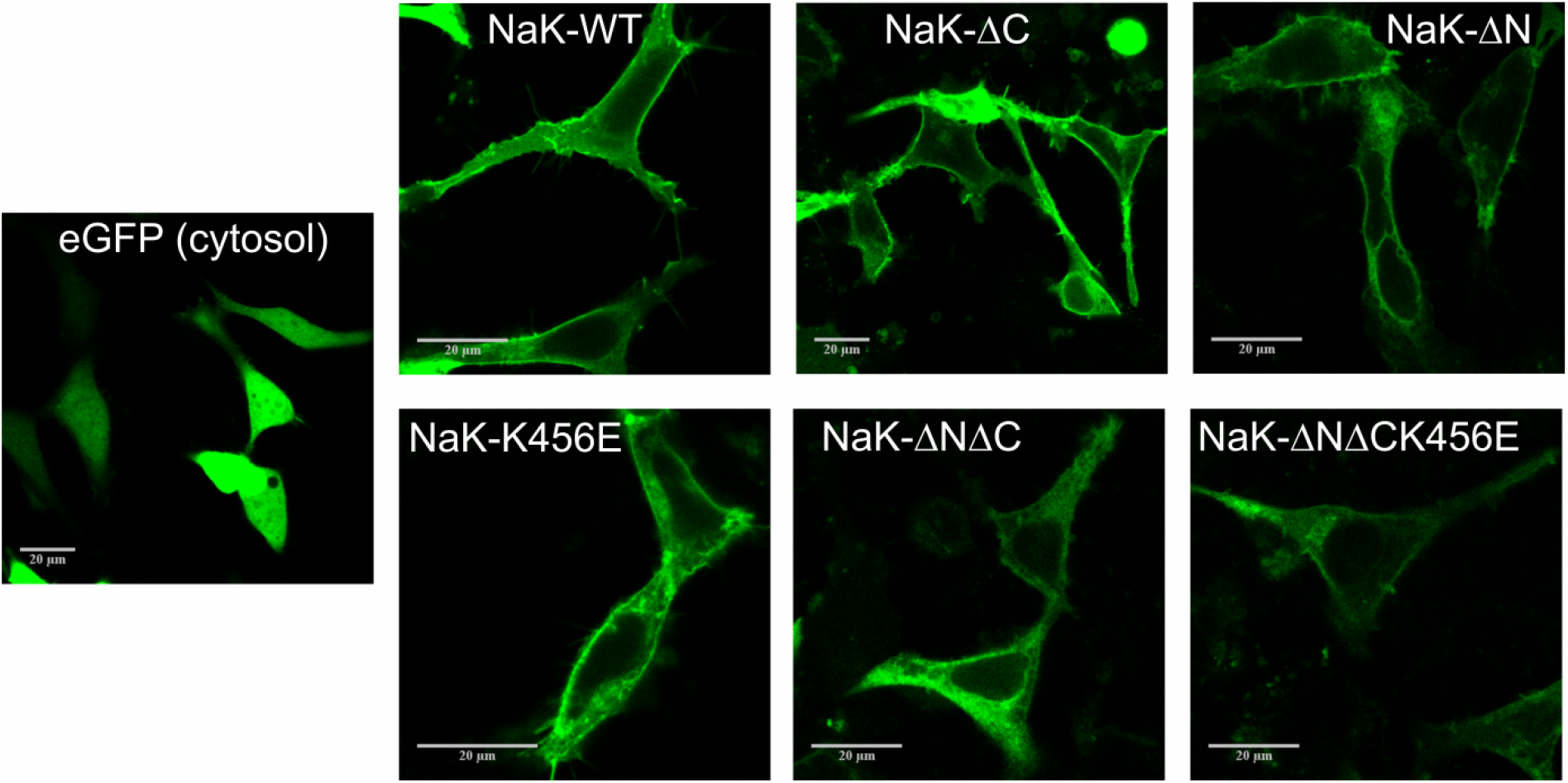
CLSM images of HEK293T cells overexpressing eGFP-labeled Na^+^,K^+^-ATPase constructs analyzed in this work. The different Na^+^,K^+^-ATPase constructs (NaK-WT, NaK-ΔN, NaK-ΔC, NaK-K456E, NaK-ΔNΔC, NaK-ΔNΔCK456E) show typical plasma membrane localization, which is different from the distribution of eGFP expressed in the cytoplasm (left). Scale bars correspond to 20 μm.

Figure 3 shows autocorrelation curves from FCS experiments of eGFP-labeled Na^+^,K^+^-ATPase constructs normalized to the same amplitude, G(τ) = 1 at τ = 0 μs. As described in previous work (Junghans et al., 2015), the eGFP-labeled wildtype Na^+^,K^+^-ATPase exhibits two diffusion times (τ_1_ = (533 ± 160) μs, amplitude fraction: (41 ± 6) %; and τ_2_ = (63 ± 18) ms, amplitude fraction: (59 ± 6) %), from which the shorter time constant is already significantly larger than the one measured for the diffusion of the isolated eGFP protein expressed in the cytoplasm ((376 ± 69) μs; (Junghans et al., 2015)).

**Figure 3:**
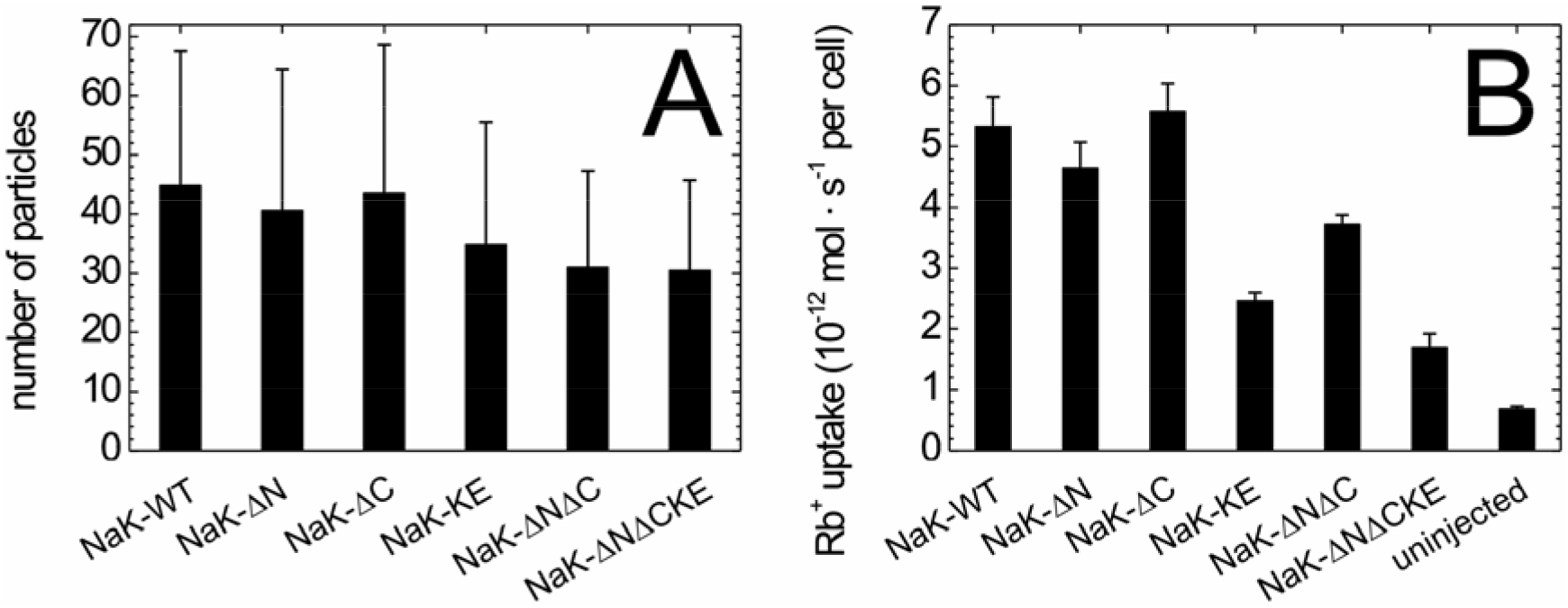
Temporal autocorrelation curves normalized to the same amplitude show differences in the lateral diffusion dynamics of different eGFP-labeled Na^+^,K^+^-ATPase constructs. (A) Comparision of NaK-WT, NaK-ΔC, NaK-ΔN and NaK-K456E, (B) comparison of NaK-WT, NaK-ΔNΔC and NaK-ΔNΔCK456E. Error bars correspond to standard deviation. In (A), error bars for each data set are shown only in one direction for clarity. Data are means ± standard deviation from 30 (NaK-WT), 31 (NaK-ΔN), 32 (NaK-ΔC), 32 (NaK-K456E), 32 (NaK-ΔNΔC) and 29 (NaK-ΔNΔCK456E) individual measurements.

Compared to NaK-WT, statistically different diffusion times and, consequently, diffusion constants were obtained for all Na^+^,K^+^-ATPase mutants, with all constructs revealing two diffusion time components (τ_1_ and τ_2_) with rather invariant relative amplitudes (Table 1). Of all single-motif disruptions, the AnkB-associated mutant (NaK-K456E) exhibited the fastest diffusion times, whereas the two Cav1-associated mutants (NaK-ΔN and -ΔC) showed an intermediate profile. Two-sample Student’s t-tests confirmed that the corresponding diffusion time components of Na^+^,K^+^-ATPase wild-type and mutants were significantly different (P < 0.05).

**Table 1:**
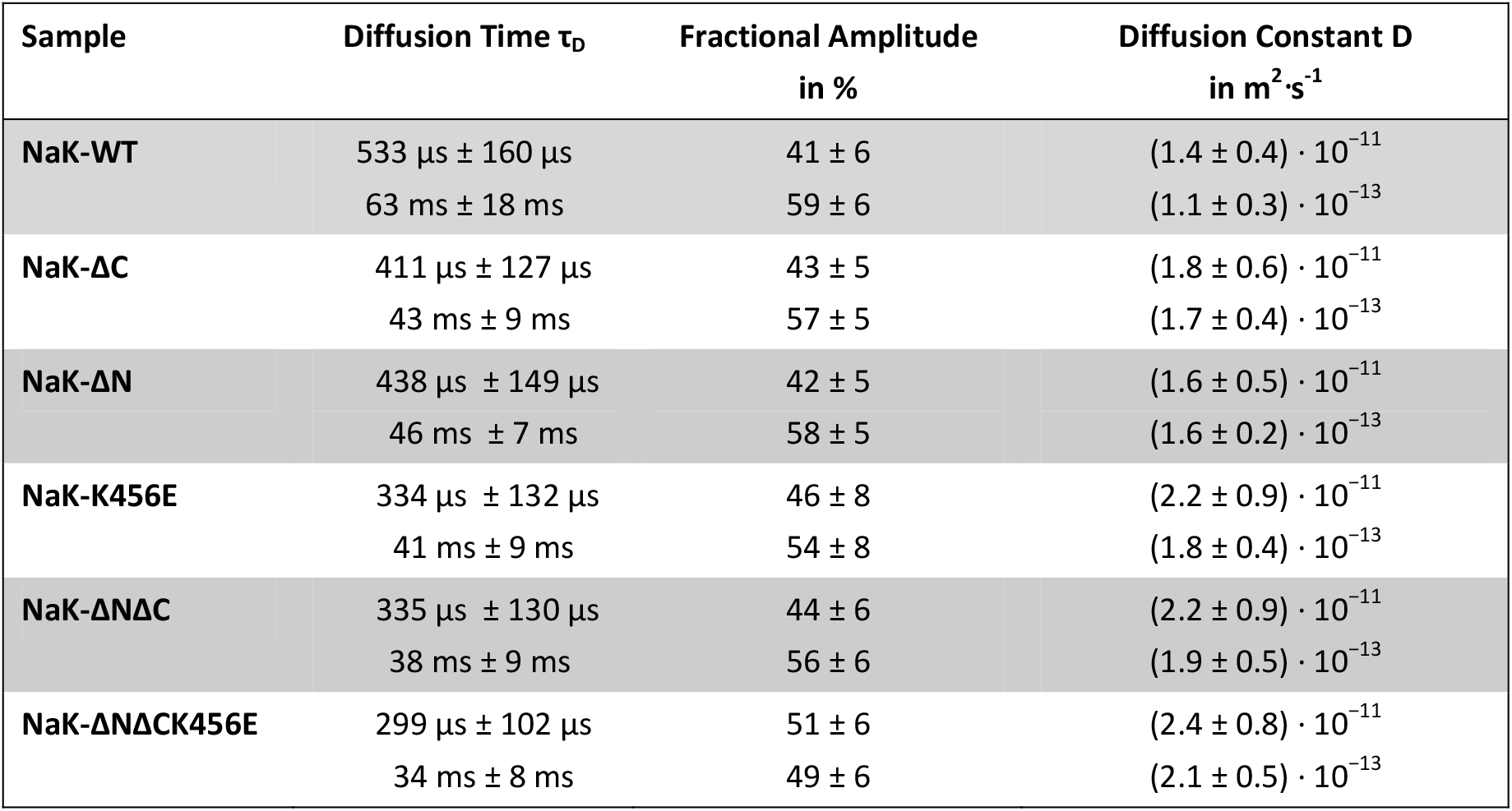
Summary of calculated correlation times with corresponding fractional amplitudes from FCS experiments and the resulting diffusion coefficients for the eGFP-labeled Na^+^,K^+^-ATPase constructs.

| Sample | Diffusion Time $\tau_D$ | Fractional Amplitude<br>in % | Diffusion Constant D<br>in $\text{m}^2 \cdot \text{s}^{-1}$ |
| --- | --- | --- | --- |
| NaK-WT | $533 \mu\text{s} \pm 160 \mu\text{s}$ | $41 \pm 6$ | $(1.4 \pm 0.4) \cdot 10^{-11}$ |
| | $63 \text{ ms} \pm 18 \text{ ms}$ | $59 \pm 6$ | $(1.1 \pm 0.3) \cdot 10^{-13}$ |
| NaK-ΔC | $411 \mu\text{s} \pm 127 \mu\text{s}$ | $43 \pm 5$ | $(1.8 \pm 0.6) \cdot 10^{-11}$ |
| | $43 \text{ ms} \pm 9 \text{ ms}$ | $57 \pm 5$ | $(1.7 \pm 0.4) \cdot 10^{-13}$ |
| NaK-ΔN | $438 \mu\text{s} \pm 149 \mu\text{s}$ | $42 \pm 5$ | $(1.6 \pm 0.5) \cdot 10^{-11}$ |
| | $46 \text{ ms} \pm 7 \text{ ms}$ | $58 \pm 5$ | $(1.6 \pm 0.2) \cdot 10^{-13}$ |
| NaK-K456E | $334 \mu\text{s} \pm 132 \mu\text{s}$ | $46 \pm 8$ | $(2.2 \pm 0.9) \cdot 10^{-11}$ |
| | $41 \text{ ms} \pm 9 \text{ ms}$ | $54 \pm 8$ | $(1.8 \pm 0.4) \cdot 10^{-13}$ |
| NaK-ΔNΔC | $335 \mu\text{s} \pm 130 \mu\text{s}$ | $44 \pm 6$ | $(2.2 \pm 0.9) \cdot 10^{-11}$ |
| | $38 \text{ ms} \pm 9 \text{ ms}$ | $56 \pm 6$ | $(1.9 \pm 0.5) \cdot 10^{-13}$ |
| NaK-ΔNΔCK456E | $299 \mu\text{s} \pm 102 \mu\text{s}$ | $51 \pm 6$ | $(2.4 \pm 0.8) \cdot 10^{-11}$ |
| | $34 \text{ ms} \pm 8 \text{ ms}$ | $49 \pm 6$ | $(2.1 \pm 0.5) \cdot 10^{-13}$ |

Upon combination of the Cav1-associated mutations in the double mutant NaK-ΔCΔN, the two diffusion times were faster than those from the corresponding single mutants and comparable to the results for NaK-K456E indicating that the mutations produce synergistic effects. The triple mutant NaK-ΔCΔNK456E exhibited the fastest diffusion times of all constructs investigated, which again indicates that Cav1- and AnkB-associated mutations act cooperatively. Two-sample Student’s t-test analysis again confirmed that the corresponding diffusion time components of Na^+^,K^+^-ATPase wildtype and double/triple mutants were significantly different (P < 0.001).

From the measured diffusion times, the corresponding diffusion coefficients were calculated according to equation 3. The calculated diffusion constants are depicted in Figure 4 and listed in Table 1 together with the diffusion times.

**Figure 4.**
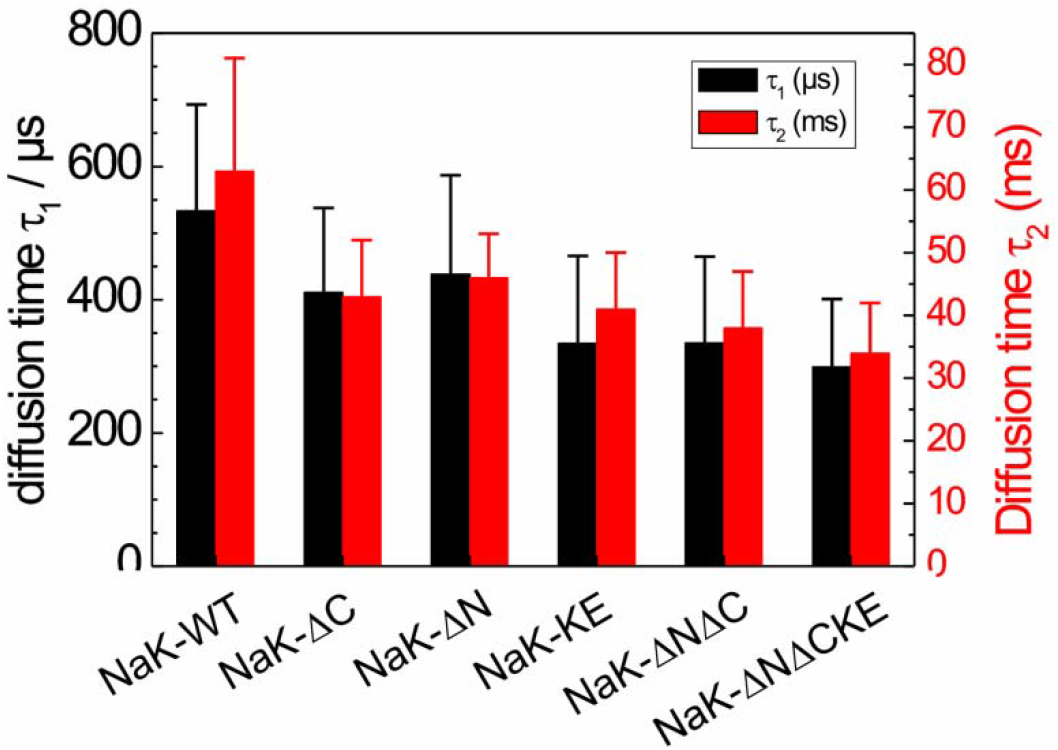
Diffusion times derived from FCS curves in Figure 3 from fits of the model function in equation 2 to the datasets. Values are means ± standard deviation.

### Activity tests by Rb^+^ uptake assays on Xenopus oocytes

Since confocal microscopy images indicated changes in the cellular distribution of some of the Na^+^,K^+^-ATPase mutants, we tested whether the density of molecules in the plasma membrane was altered by the mutations. Since the G(0) value, which represents the amplitude of the autocorrelation curve at time zero, corresponds to the reciprocal of the number of molecules in the observation volume element (OVE), it can be used as a measure of the density of molecules in the plasma membrane. Figure 4A shows the average numbers of molecules in the OVE for the Na^+^,K^+^-ATPase constructs tested. Despite large error bars, the data by trend indicate that the NaK-K456E mutant, and even more so the double (NaK-ΔNΔC) and triple mutant (NaK-ΔNΔCK456E) are less abundant in the plasma membrane than the wildtype enzyme. To further test the functional effects of mutations, we quantified the ion-pumping function of the mutated Na^+^,K^+^-ATPase constructs.

To this end, the eGFP-tagged human Na^+^,K^+^-ATPase α_2_-subunit constructs were expressed (together with human Na^+^,K^+^-ATPase β_1_-subunit) in *Xenopus* oocytes, and pumping activity was examined by Rb^+^ uptake measurements. After loading the cells with Na^+^ to activate the ion pump (see Methods), cells were incubated in solution containing 1 mM RbCl for three minutes. Subsequently, the Rb^+^ content of individual cells was analyzed by atomic absorption spectrophotometry (AAS) using a transversely heated graphite tube furnace for atomization. Figure 5B shows Rb^+^ uptake values for the various Na^+^,K^+^-ATPase mutants in comparison to the wildtype enzyme, whereas uninjected oocytes served as control for non-specific Rb^+^ uptake. Typical Rb^+^ uptake values were about 5 pmoles (5·10^−12^ moles) per second per cell, which, based on a turnover number of human α_2_/β_1_ Na^+^,K^+^-ATPase of about 13 s^−1^ (Tavraz et al., 2008) at 22 °C, corresponds to an average of 1.2·10^11^ pump molecules in the plasma membrane of *Xenopus* oocytes. In terms of pump current (due to the 3 Na^+^: 2 K^+^ transport stoichiometry, one net transported charge corresponds to two transported Rb^+^ ions), 5 pmoles of Rb^+^ per second per oocyte corresponds to about 200 nA pump current, which is a typical value for electrophysiological experiments on Na^+^,K^+^-ATPase in *Xenopus* oocytes (Tavraz et al., 2008) indicating that the eGFP label at the N-terminus does not alter functional activity of the wildtype enzyme. However, some of the mutants apparently exhibited lower activity. The data in Figure 5B show that the Cav1-associated mutants NaK-ΔN and -ΔC had about the same transport activity as the wildtype enzyme, but Rb^+^ flux was substantially reduced for the NaK-K456E construct. The double mutant NaK-ΔNΔC showed slightly reduced Rb^+^ uptake compared to wildtype, and the triple mutant NaK-ΔNΔCK456E exhibited the smallest uptake activity of all constructs tested. These observations indicate that the reduced plasma membrane localization of the NaK-K456E, NaK-ΔNΔC and NaK-ΔNΔCK456E variants seen in CLSM images correlates, by trend, with reduced molecule density in the plasma membrane. In mutants with disrupted AnkB interaction motif (NaK-K456E, NaK-ΔNΔCK456E) the reduced transport to the plasma membrane is well reflected by reduced overall pumping activity.

**Figure 5.**
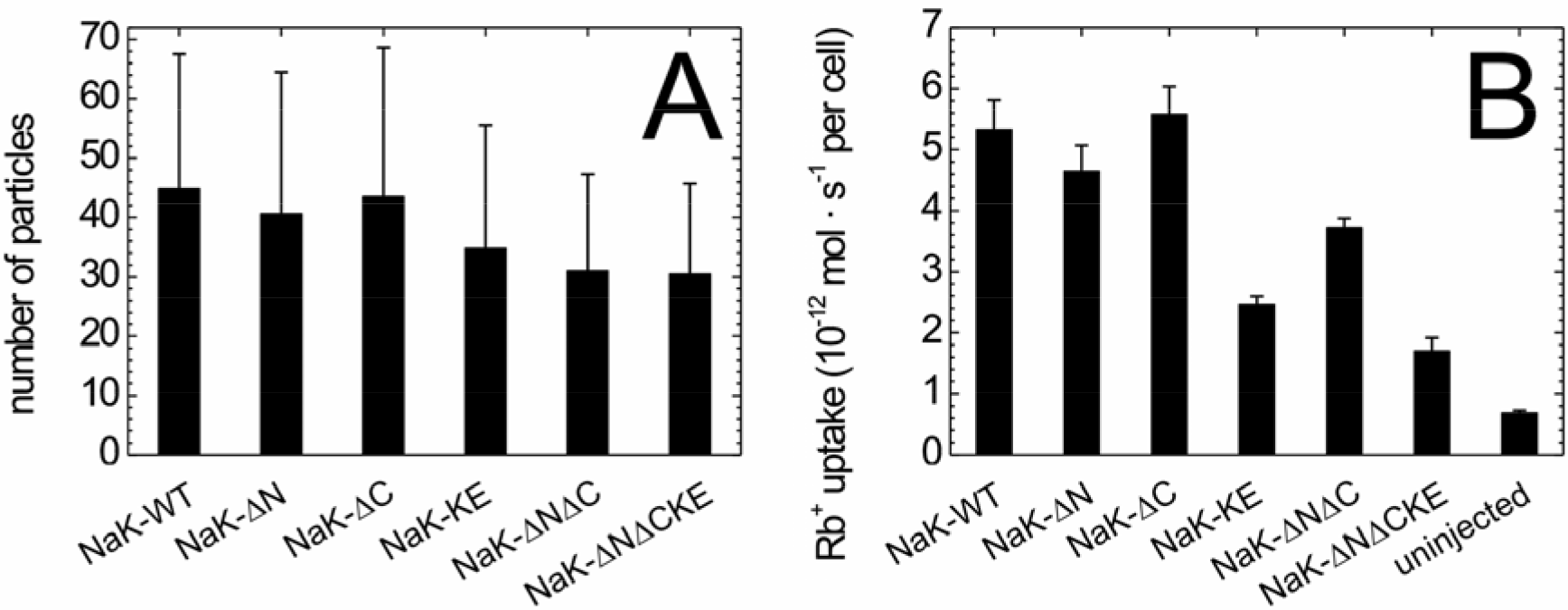
Average number of molecules in the OVE and Rb+ uptake activity for different Na^+^,K^+^-ATPase constructs. (A) From the G(0) values of the autocorrelation curves (Figure 3), the average numbers of molecules in the observation volume element were calculated. Data are means ± standard deviation from 30 (NaK-WT), 29 (NaK-ΔN), 31 (NaK-ΔC), 30 (NaK-K456E), 31 (NaK-ΔNΔC) and 29 (NaK-ΔNΔCK456E) individual measurements. (B) Rb^+^ uptake activity of the various eGFP-labeled Na^+^,K^+^-ATPase constructs expressed in *Xenopus* oocytes as determined by atomic absorption spectrophotometry. Data are means ± standard errors, for dataset, between 14 and 30 oocytes from two or three independent batches were measured.

### FRAP and FRAS experiments

Another method to characterize the diffusion behavior of molecules in membranes is fluorescence recovery after photobleaching (FRAP). To utilize fluorescence recovery as readout, we employed two experimental schemes. First, we used the standard method, in which fluorescence-labeled Na^+^,K^+^-ATPase molecules were bleached with intense confocal scanning illumination, which we refer to herein as “irreversible” bleaching. For this procedure, we used Na^+^,K^+^-ATPase constructs labeled with eGFP (as used in FCS experiments), or with the photoswitchable derivative Dreiklang (DRK), which was the first fluorescent protein variant, for which the wavelength ranges for ON- and OFF-switching as well as for fluorescence excitation are well separated (Brakemann et al., 2011). The second employed technique relied on N-terminally DRK-labeled Na^+^,K^+^-ATPase constructs, since the DRK fluorophore permits reversible OFF-switching by low light intensities instead of irreversible bleaching. In principle, such an OFF-switching experiment with DRK could be repeated multiple times on the same cell membrane. Therefore, we refer to this scheme as “reversible” bleaching or fluorescence recovery after photoswitching (FRAS). Figure 6 shows a series of confocal fluorescence images from a FRAP experiment using eGFP-labeled Na^+^,K^+^-ATPase.

**Figure 6:**
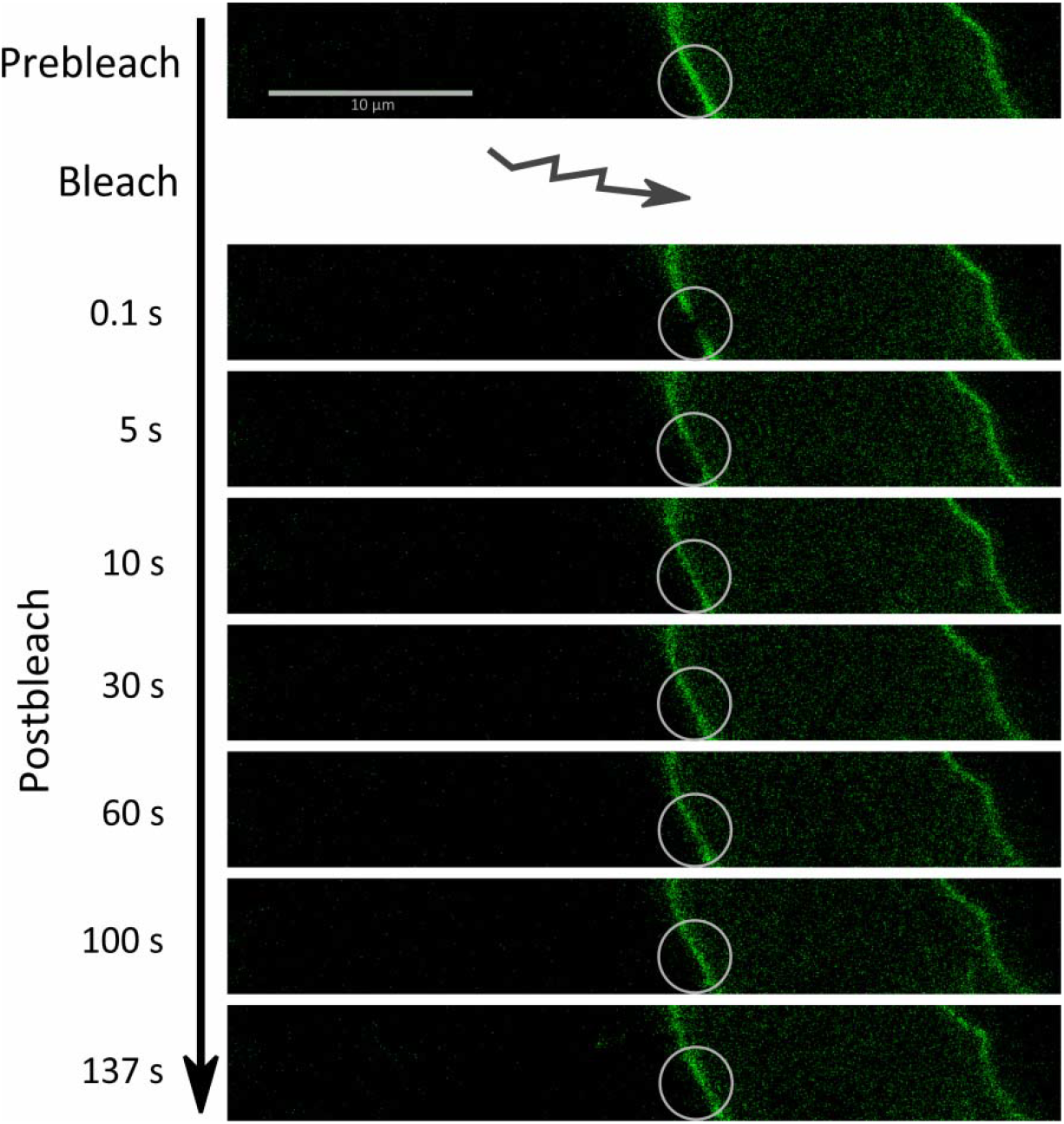
Exemplary FRAP time series from a HEK293T cell expressing eGFP-labeled wildtype Na^+^,K^+^-ATPase. White circles indicate the region of interest (ROI) before bleaching (top) and at various times after bleaching, as indicated on the left.

Average normalized fluorescence recovery curves for fluorescence-labeled Na^+^,K^+^-ATPase wildtype and the Cav1- and AnkB-related mutants are shown in Figure 7. Notably, the recovery curves acquired with the reversible FRAS scheme (Figure 7A) were characterized by much longer recovery times (t_1_/_2_ values from exponential fits with two characteristic times, see Table 2) than the curves measured based on irreversible FRAP, irrespective of the fluorophore used (Figure 7B,C). This is even more striking, since recovery in irreversible FRAP should only contain contributions from nonbleached molecules diffusing into the pre-bleached area, whereas the reversible FRAS scheme should contain an additional contribution from slow thermal recovery of OFF-switched DRK molecules into the fluorescent ON state. Moreover, only in FRAS experiments, the recovery times of the Na^+^,K^+^-ATPase mutants were significantly (P<0.05, Student’s t-test) longer than the one of the wildtype enzyme, with the same trend among the mutants as observed in FCS experiments. In contrast, the much faster FRAP recovery curves did not show any differences between wildtype and mutants. The abnormally fast recovery curves from irreversible FRAP point at a common disadvantage of the proteinaceous GFP/DRK chromophores, which exhibit a much more complex photophysical behavior than smaller organic dye molecules, with population of dark-states, triplet transitions, blinking etc. Such problems in utilizing FPs as fluorescence tags for FRAP experiments have been described in the literature before (Sinnecker et al., 2005) indicating that the irreversible FRAP curves rather reflect complex photophysical transitions from multiple “dark” states to the fluorescent state, which precludes the observation of mutation-related differences in diffusion behavior.

**Figure 7:**
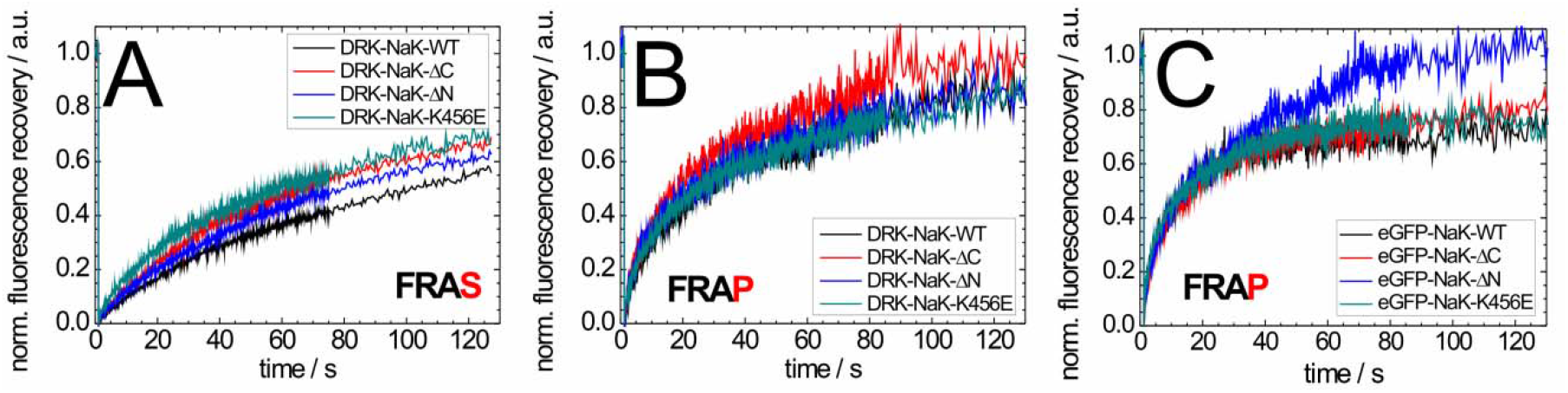
FRAS and FRAP experiments with Na^+^,K^+^-ATPase constructs N-terminally labeled with the photoswitchable DRK (A,B) or eGFP (C) upon expression in HEK293T cells. (A) “Reversible” FRAS scheme using low-intensity OFF-switching of DRK by illumination with 405 nm light. Excitation used 514 nm laser light, fluorescence emission was detected the 520-600 nm range. Data are averages from 19 (NaK-WT), 19 (NaK-ΔN), 18 (NaK-ΔC) and 19 (NaK-K456E) individual measurements. (B) “Irreversible” FRAP scheme using intense illumination with 514 nm laser light to bleach DRK fluorescence. Data are averages from 16 (NaK-WT), 20 (NaK-ΔN), 20 (NaK-ΔC) and 24 (NaK-K456E) individual measurements. (C) “Irreversible” FRAP scheme using intense illumination with 488 nm laser light to irreversibly bleach eGFP fluorescence. Data are averages from 20 (NaK-WT), 18 (NaK-ΔN), 19 (NaK-ΔC) and 20 (NaK-K456E) individual measurements.

**Table 2:** Summary of determined mobile fractions (M), t_1_/_2_-values and calculated diffusion constants *D* for different Na^+^,K^+^-ATPase constructs from FRAS and FRAP experiments. t_1_/_2_ represents the time of half-maximal recovery of the normalized fluorescence intensity determined from fitting the curves in Figure 7 with Equation 6. Mobile fractions *M* were determined according to Equation 8 from the fitted curves and diffusion constants *D* according to Equation 7. All values are means ± standard deviation from the number of measurements given in the legend of Figure 7.

| Sample | Method | Mobile Fraction $M$<br>% | $t_{1/2}$ in s | Diffusion Constant<br>$D$ in $\text{m}^2/\text{s}$ |
| --- | --- | --- | --- | --- |
| DRK-NaK-WT | reversible<br>(FRAS) | $87 \pm 17$ | $78 \pm 37$ | $(3.2 \pm 1.5) \cdot 10^{-15}$ |
| DRK-NaK-$\Delta\text{C}$ | | $83 \pm 22$ | $46 \pm 21$ | $(5.4 \pm 2.5) \cdot 10^{-15}$ |
| DRK-NaK-$\Delta\text{N}$ | | $81 \pm 19$ | $52 \pm 23$ | $(4.8 \pm 2.1) \cdot 10^{-15}$ |
| DRK-NaK-K456E | | $81 \pm 24$ | $39 \pm 23$ | $(6.4 \pm 3.8) \cdot 10^{-15}$ |
| DRK-NaK-WT | Irreversible<br>(FRAP) | $85 \pm 18$ | $19 \pm 12$ | $(1.3 \pm 0.8) \cdot 10^{-14}$ |
| eGFP-NaK-WT | | $75 \pm 22$ | $8 \pm 6$ | $(3.1 \pm 2.3) \cdot 10^{-14}$ |
| DRK-NaK-$\Delta\text{C}$ | | $94 \pm 13$ | $17 \pm 9$ | $(1.5 \pm 0.8) \cdot 10^{-14}$ |
| eGFP-NaK-$\Delta\text{C}$ | | $82 \pm 19$ | $11 \pm 8$ | $(2.3 \pm 1.7) \cdot 10^{-14}$ |
| DRK-NaK-$\Delta\text{N}$ | | $87 \pm 15$ | $18 \pm 11$ | $(1.4 \pm 0.9) \cdot 10^{-14}$ |
| eGFP-NaK-$\Delta\text{N}$ | | $93 \pm 11$ | $12 \pm 5$ | $(2.1 \pm 0.9) \cdot 10^{-14}$ |
| DRK-NaK-K456E | | $85 \pm 17$ | $18 \pm 12$ | $(1.4 \pm 0.9) \cdot 10^{-14}$ |
| eGFP-NaK-K456E | | $79 \pm 19$ | $8 \pm 6$ | $(3.1 \pm 2.3) \cdot 10^{-14}$ |

Using DRK as fluorescence marker can potentially avoid artifacts induced by high-intensity laser illumination, but, it needs to be said that the observed recovery curves on these time scales also contain (unknown) contributions from thermal conversion from the OFF to the ON state, which would have to be determined from more detailed *in vitro* and *in vivo* experiments on (purified) DRK protein. However, the FRAS scheme utilizing the DRK label indeed allows discriminating differences in the diffusion behavior of Na^+^/K^+^-ATPase mutants, which are qualitatively similar to FCS, and the possibility to repeat the reversible photoswitching scheme would allow accumulating more data from the same set of cells to increase the significance level substantially. As another observation, we note that the diffusion constants determined from FRAP and FRAS experiments differ by about one or even two orders of magnitude, respectively, from the values obtained from FCS experiments. Such discrepancies between these two seemingly similar experimental techniques have frequently been reported in the literature, even if applied to the same molecular objects. For example, for G-protein coupled receptors, diffusion coefficients in the order of 10^−13^ m^2^·s^−1^ were determined by FCS and 10^−15^ m^2^·s^−1^ by FRAP (Calizo and Scarlata, 2013). Similar effects were observed for a YFP-labeled dopamine transporter or eGFP-labeled epidermal growth factor receptor and β-adrenergic receptor (Adkins et al., 2007). The authors of the latter study also showed that the resulting diffusion coefficients depend on the used cell line.

## Discussion

In this work, the dependence of the mobility of the Na^+^,K^+^-ATPase α_2_-subunit in the plasma membrane of HEK293T cells on the disruption of known binding motifs for Cav1 and/or AnkB was probed by FCS and FRAP/FRAS. Since the Na^+^,K^+^-ATPase studied here is a single entity composed of an α_2_- and a β_1_-subunit, one would expect only a single characteristic diffusion time. But, as outlined in previous work (Junghans et al., 2015), the autocorrelation curves already for the wildtype Na^+^,K^+^-ATPase are complex, with two diffusion components (τ_1_ ≈ 500 μs and τ_1_ ≈ 60 ms) separated by two orders of magnitude corresponding to diffusion coefficients of 1.4 · 10^−11^ m^2^ ·s^−1^ and 1.1 · 10 m^−13^ m^2^·s^−1^, respectively, indicating the presence of two independent Na^+^,K^+^-ATPase populations with largely different mobility. Such multi-component autocorrelation curves are not uncommon for membrane proteins (Chiantia et al., 2009; Schwille et al., 1999b; Vukojevic et al., 2008) and reflect the complexity of the membrane environment that allows for interactions with certain lipid domains, other membrane proteins, the cytoskeleton or other scaffolding partners, but also exclusion from some domains. However, rationalization of either of the two diffusion components is all but straightforward.

In the literature, the diffusion time is commonly estimated from the molecular weight of a protein. The eGFP-labeled Na^+^,K^+^-ATPase α_2_/β_1_-complex has a molecular weight of about 183.5 kDa, which would result in a diffusion time of around 1.8 ms across the size of the OVE. From a rough approximation treating the diffusing molecules as spheres with hydrodynamic radii proportional to the cubic root of the molecular weight (eGFP: 26.9 kDa, eGFP-labeled Na^+^,K^+^-ATPase α-subunit plus glycosylated β-subunit: 183.5 kDa) the diffusion coefficient should increase by a factor of 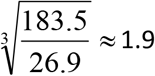 for the change from isolated eGFP to eGFP-labeled Na^+^,K^+^-ATPase. Of course, this estimation neglects (i) that the actual shape of the molecules is different from a sphere, (ii) that the Na^+^,K^+^-ATPase can diffuse only in two dimensions, and (iii) that the dynamic viscosity of biological membranes is different from that of the cytoplasm. The fast diffusion coefficient of the eGFP-labeled Na^+^,K^+^-ATPase in the plasma membrane (1.4 · 10^−11^ m^2^·s^−1^) is only about 1.3-fold smaller than the diffusion time of eGFP in the cytoplasm (*D_eGFP_* = 1.9 · 10 ^−11^ m^2^·s^−1^, (Junghans et al., 2015)), which still agrees with the above estimation within error limits suggesting that the fast-diffusing fraction of Na^+^,K^+^-ATPase molecules diffuses rather freely. However, the slow diffusion component cannot be rationalized, even if one assumes that interactions of Na^+^,K^+^-ATPase with Cav1 macrostructures composed of 14 to 16 monomers (molecular weight between 300 kDa up and 400 kDa (Williams and Lisanti 2004)) are possible. Thus, formation of large hetero-oligomeric protein complexes cannot explain the extremely long diffusion times (Elson, 2001) and anchoring of Na^+^,K^+^-ATPase molecules to components of the cytoskeleton or trapping in certain membrane nanodomains needs to be taken into account.

Literature data on Na^+^/K^+^-ATPase for a direct comparison is sparse and essentially stems from FRAP studies carried out decades ago. Paller and coworkers studied Na^+^,K^+^-ATPase in renal proximal tubule epithelial cells upon labeling with 9-anthroylouabain, a fluorescent derivative of the specific inhibitor of the Na^+^ pump, ouabain, and reported a diffusion constant of 3.3 · 10^−14^ m^2^·s^−1^ (at 25 °C), which increased to 2.4 · 10^−13^ m^2^·s^−1^ upon treatment with cytochalasin A, an agent causing actin depolymerization and disruption of the cytoskeleton (Paller, 1994). Vaz and coworkers studied the related Ca^2+^-ATPase (about 100 kDa) from rabbit muscle sarcoplasmic vesicles labeled with fluorescein iodoacetamide and reconstituted in liquid-crystalline phase bilayers. For this system, diffusion coefficients of 1.8 · 10^−12^ m^2^·s^−1^ (at 36 °C) and 9.9 · 10^−13^ m^2^·s^−1^ (at 13 °C) were found (Vaz et al., 1982). By order of magnitude (10^−13^ m^2^·s^−1^), these literature FRAP data correspond to the slow diffusion constants determined by our FCS approach, but are still two orders of magnitude faster than the parameters determined by the reversible FRAS scheme. It should be noted that membrane protein diffusion coefficients can vary largely dependent on the cell line used, as well as for different measurement techniques (Feder et al., 1996). For other membrane-embedded proteins, recent publications list values between 10^−16^ cm^2^·s^−1^ and 10^−14^ cm^2^·s^−1^ (Kühn et al., 2011), and a more recent FRAP study reported values between 2 · 10^−14^ cm^2^·s^−1^ and 2.2 · 10^−13^ cm^2^·s^−1^ depending on the number of transmembrane segments (Kumar et al., 2010). However, such slow diffusion constants have not only been found for membrane proteins. Even for a farnesyl-anchored eGFP, the study by Ohsugi et al. reported a lateral diffusion coefficient of 5.6 · 10^−13^ cm^2^·s^−1^ in the plasma membrane of COS-7 cells showing that even a fluorescent protein with a low-molecular-weight membrane anchor can exhibit significantly retarded diffusion, which is by two orders of magnitude slower than free diffusion in the cytoplasm (Ohsugi et al., 2006).

In theory, FCS and FRAP should yield comparable information, however, there are practical differences between FCS and FRAP and these two methods often yield different values for the diffusion coefficient of plasma membrane associated proteins. Whereas FCS detects the diffusion of individual molecules through an OVE of diffraction-limited size, FRAP monitors the diffusion of large ensembles of molecules into a comparatively large pre-bleached membrane area that is several μm in diameter. In addition, immobile molecules are invisible in FCS, since they don’t give rise to fluorescence intensity fluctuations, but rather contribute to the background. Moreover, slowly moving molecules may be photobleached before they exit the observation volume element, giving rise to shorter decay times that can be mistaken for faster diffusion. Therefore, FCS systematically overestimates the diffusion coefficient of investigated molecules. In comparison, FRAP is biased by the fact that photobleaching is not instantaneous; as photobleaching is performed, molecules enter/exit the observation volume element and the photobleached area is typically larger than the area that is specified by the region of interest (ROI). As a consequence, a large pool of molecules is photobleached. In addition, subsequent image acquisition may suppress the fluorescence signal recovery. Hence, FRAP often yields diffusion coefficient values that are systematically underestimated.

### Theoretical considerations and models of membrane diffusion

The ‘fluid mosaic model’ (Singer and Nicolson, 1972) founded the modern concept of the plasma membrane as a two-dimensional liquid, in which all membrane-embedded proteins and lipids diffuse freely. Saffman and Delbrück developed a membrane model for two-dimensional diffusion in 1975 (Saffman and Delbrück, 1975), which considers membrane proteins as cylindrical inclusions of radius *R*_0_ that diffuse in an infinite (and otherwise homogenous) bilayer of thickness *h* and viscosity *η_m_* within a solvent of viscosity *η_s_*. The Saffman-Delbrück model results in a weak, logarithmic dependence of the diffusion constant *D* on the particle radius (here, *γ* ≈ 0,5772 is the Euler-Mascheroni constant):

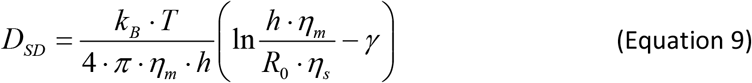

A recent dual-focus FCS study on the diffusion dynamics of membrane proteins varying largely in size (11.3 kDa / 0.7 nm radius for cytochrome b5, and 345 kDa / 3.6 nm radius for a trimeric ABC transporter, AcrB) reconstituted in artificial black lipid membranes reported diffusion coefficients between 8.5 · 10^−12^ m^2^·s^−1^ and 10.2 · 10^−12^ m^2^·s^−1^ (Weiss et al., 2013), in compliance with the Saffman-Delbrück model (Figure 8). However, other authors found a stronger 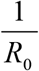 dependence of *D* on the radius resembling Stokes-Einstein behavior (Gambin et al., 2006):

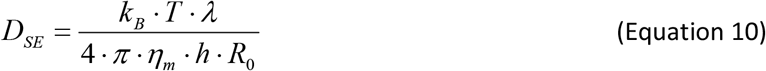

**Figure 8:**
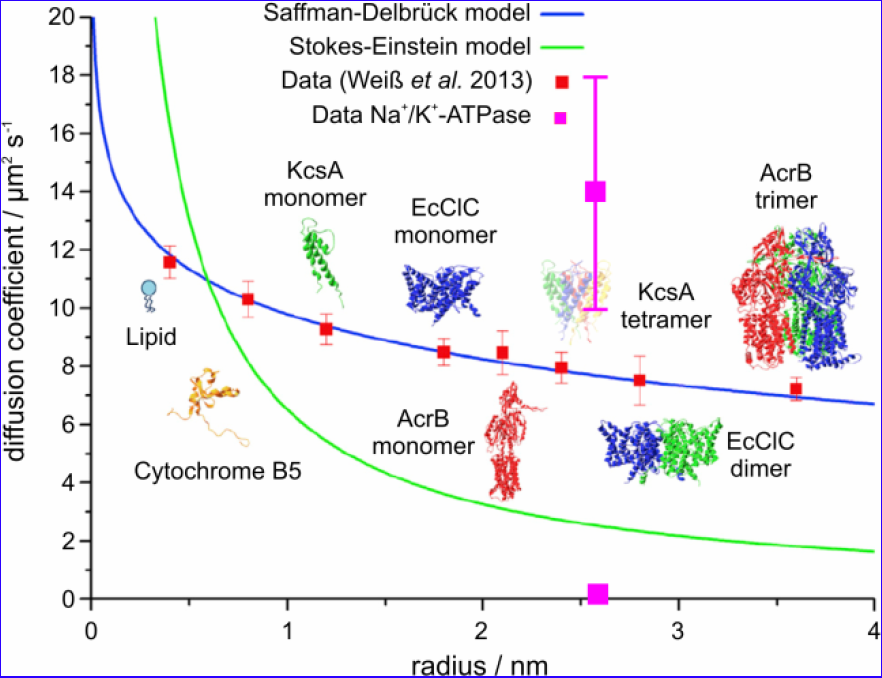
Comparison of the obtained diffusion constants of Na^+^,K^+^-ATPase (magenta) to literature data (graph redrawn with permission from (Weiss et al., 2013)), in which diffusion of several membrane proteins of largely varying size was measured in artificial lipid bilayers by dual-focus FCS. The fast diffusion constant of the Na^+^,K^+^-ATPase is much larger than those of the model proteins studied by Weiß and coworkers, which by themselves complied well with the Saffman-Delbrück model. The radius of the transmembrane part of the Na^+^/K^+^-ATPase α/β-complex was estimated from the crystal structure PDB 3A3Y (Shinoda et al., 2009).

For dimensional reasons, the parameter *λ* is introduced in Equation 10 as a characteristic length scale. Justifications for the Stokes-Einstein behavior were given based on e.g. hydrophobic mismatch or changes in bulk hydrodynamics due to height mismatch between the membrane and the embedded protein (Naji et al., 2007). The question whether Saffman-Delbrück or Stokes-Einstein behavior governs diffusion of membrane proteins cannot be resolved here and is still a matter of debate. However, it should be noted that the Na^+^,K^+^-ATPase’s diffusion constant of the fast process determined here is even too large, i.e., diffusion is apparently “too fast” compared to the model proteins studied in artificial membranes by Weiß *et al*. (Figure 8) showing that artificial bilayers cannot serve as a good reference for diffusion processes in biological membranes.

### Super- and subdiffusion phenomena in membranes

Numerous publications suggest that membrane proteins are prone to exhibit anomalous diffusion instead of normal Brownian diffusion. In case of anomalous diffusion, the mean-square displacement (MSD) of molecules no longer scales linearly with the diffusion time τ_*D*_, as in the case of free Brownian motion (Equation 3). The MSD is rather proportional to 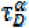, with subdiffusion characterized by 0 < *α* ≤ 1 and superdiffusion by *α* > 1. Only for *α* = 1, the molecules diffuse normally (Chiantia et al., 2009; Elson, 2001; Feder et al., 1996; Schwille et al., 1999a). The complexity of the plasma membrane, with its composition of various lipids or cholesterol, with membrane proteins embedded that may or may not form complexes with others or may even form linkages to cytosolic and cytoskeletal proteins, creates a structural organization, which is characterized by a hierarchy of different length scales. Due to this hierarchical patterning, also the apparent properties of diffusion phenomena depend on the length scale, across which diffusion is actually monitored. Along these lines, Wawrezinieck *et al*. suggested that the diffusion behavior of membrane proteins must be considered as a function of the size of the OVE. FCS measurements in the plasma membrane probed for one size of OVE are difficult to interpret in the presence of membrane microdomains and linkages to the cytoskeleton meshwork, especially if the diffusion behavior of largely different membrane components such as lipids or transmembrane proteins is evaluated (Wawrezinieck et al., 2005). On the small scale, the lipid composition gives rise to particular microdomains such as lipid rafts, in which cholesterol is tightly packed with long saturated fatty acid chains of glycosphingolipids, which in effect favors phase separation processes in the lipid bilayer to create microdomains differing largely in fluidity. On the long scale, interactions with the cellular cytoskeleton and other cellular matrix proteins, as well as integral membrane proteins lead to the formation of a heterogeneous protein meshwork. Therefore, Wawreziniek, Lenné and coworkers defined diffusion laws, which describe diffusion on different length scales, resulting in four different diffusion models for the membrane organization to simulate these diffusion laws. When tested regarding the existence of diffusion barriers and microdomains within the plasma membrane of COS-7 cells (Lenné et al., 2006), the study of Lenné *et al*. could confirm the applicability of the hierarchical method of analysis.

Two other models have been also proposed in the literature, the membrane-skeleton "fence" model (also termed membrane-skeleton "corralling" model) and the anchored protein "picket" model, which both can be viewed as variants of the “meshwork” model. The former suggests that a fence is built up at the intracellular interface of the lipid bilayer by the actin filament network and leads, therefore, to small compartments. Transmembrane proteins are thus hindered in diffusion, which is then resulting in shortened diffusion times within one compartment. Kusumi *et al*. also assumed that the transmembrane proteins can hop between the different compartments, which lead to a longterm hop diffusion for transmembrane proteins. The anchored-protein "picket" model works for all membrane proteins and assumes that different transmembrane proteins are anchored to the membrane-skeleton fence and lined up in there (Kusumi et al., 2005; Kusumi et al., 2010). In terms of the Na^+^,K^+^-ATPase studied here, the very long diffusion time component can be attributed to the dynamic patterning of biological membranes by linkage to cytoskeletal “meshworks”, “fences” or anchoring “pickets”, to which also the studied interactions with caveolin-1 and ankyrin B might well contribute.

The fast diffusion coefficient of the Na^+^,K^+^-ATPase apparently hints at superdiffusion. However, this could have trivial reasons to start with. FCS measurements on the plasma membrane always contain signal contributions from the cytoplasm, which exhibit shorter diffusion times (Ohsugi et al., 2006; Schwille et al., 1999a), because in a conventional confocal setup, the length of the OVE in axial direction is substantially larger than its radius in the plane perpendicular to the optical axis. More sophisticated explanations relate to an apparently reduced size of the observation volume element based on the microdomain model assuming that certain membrane domains are inaccessible for the diffusing species and, thus, cause artificially short diffusion times. One of the biggest challenges of FCS on plasma membranes is that the “effective” area that is available for diffusion is unknown. It is always assumed that the total OVE area is available for diffusion, and diffusion coefficients are calculated based on this assumption. However, such “excluded spaces” could be large and include cytoskeleton-anchored complexes of membrane proteins, or rigid microdomains such as lipid-rafts. Also, it needs to be mentioned that in a FCS experiment, all locally fixed fluorophores will be bleached, thus completely immobile Na^+^,K^+^-ATPase molecules carrying the eGFP label will not show up in the observed fluorescence fluctuation traces, but also create inaccessible spaces that do not allow other molecules to enter. Also, the increased affinity of certain membrane proteins to special lipids may lead to accumulation in particular microdomains (Chiantia et al., 2009), which also biases diffusion of membrane proteins because either they cannot escape from certain microdomains (hence, apparently diffusing slowly) or do not have access to it. Such considerations might especially apply for caveloae, which are special lipid rafts with high cholesterol content forming invaginations of the plasma membrane (Williams and Lisanti, 2004). Caveolae are characterized by the presence of Cav1, which is a scaffolding membrane protein for which interactions with Na^+^,K^+^-ATPase have been extensively discussed in the literature. Cai *et al*. showed by FRAP measurements on LLCPK1 cells that Cav1 is highly immobile within the plasma membrane (Cai et al., 2008). Xie *et al*. assumed that enzymes like the Na^+^,K^+^-ATPase and other signaling proteins concentrate in caveolae, which might then lead to the formation of large signaling complexes (Xie and Cai, 2003), which trap a certain fraction of Na^+^,K^+^-ATPase molecules.

### Conclusions

In summary, our FCS and FRAS studies show that mutations in all previously described interaction motifs of the Na^+^,K^+^-ATPase with Cav1 and AnkB alter the lateral mobility of the enzyme in mammalian cell plasma membranes, and that the detected changes affect both of the observed diffusion times similarly, which indicates that disruption of the interactions are effective even on different levels of spatial organization of the plasma membrane.

Second, the effects of the individual interaction motif mutations are additive, and this synergy of effects points at binding sites which are utilized or operate independent from each other.

Moreover, the mutation in the AnkB binding site also affects the plasma membrane expression level, in line with the established role of AnkB in plasma membrane targeting of the enzyme. Liu *et al*. studied eGFP-labeled Na^+^,K^+^-ATPase in plasma membrane signaling microdomains of COS-7 cells, in which Na^+^,K^+^-ATPase, IP_3_R and AnkB are active partners (Liu et al., 2008). Stabach *et al*. showed that ankyrin facilitates the passage of Na^+^,K^+^-ATPase through the secretory pathway in polarized cells, indicating that ankyrin is not only present on the plasma membrane but also in the early secretory pathway (Stabach et al., 2008), which rationalizes the observed effects on cellular distribution, plasma membrane density and functional activity.

Our complementary FRAP experiments showed that the use of genetically encoded chromophores is problematic due to their complex photophysics. However, the novel FRAS scheme, which exploits the superior properties of the reversibly photoswitchable DRK and works at much lower excitation energies, at least qualitatively discloses similar effects of the mutations, although the individual diffusion constants determined by FCS and FRAP/FRAS differ by one order of magnitude, which can, in part, be due to systematically inherent differences of the used techniques.

Our results set the stage for addressing the effects of disease-related mutations of Na^+^,K^+^-ATPase isoforms on the level of protein dynamics and their involvement in signaling complexes or specialized membrane microdomains performing highly organized signaling functions. Future studies should focus on the mobility of Na^+^,K^+^-ATPase mutants, which are associated with human inherited diseases especially if other functional tests were inconclusive or contradictory. Differences in diffusion behavior could indicate, whether some FHM, RDP or AHC mutations are associated with altered interactions to Na^+^,K^+^-ATPase binding partners. Eventually, such studies might help to establish a more comprehensive understanding of the pathophysiology of these diseases on the cellular level.

## Acknowledgements

The authors thank Agneta Gunnar (KI Stockholm) for technical support, Dr. Marco Vitali and Prof. Rudolf Rigler for stimulating discussions to initiate the work, Dr. Susan Spiller for help with *Xenopus* oocyte expression for Rb^+^ flux measurements, and Dr. Oliver Kobler (Leibnitz Institute of Neurobiology, Magdeburg) for instructions on FRAP experiments. This work was supported by travel grants for short-term scientific missions within the COST MP1205 framework (C. J.), the Swedish Research Council (V. V.) and the German Research Foundation, Cluster of Excellence “Unifying Concepts in Catalysis” (T. F.).

